# GRAIGH: Gene Regulation accessibility integrating GeneHancer database

**DOI:** 10.1101/2023.10.24.563720

**Authors:** Lorenzo Martini, Alessandro Savino, Roberta Bardini, Stefano Di Carlo

## Abstract

Single-cell assays for transposase-accessible chromatin sequencing data are one of the most powerful tools for studying the epigenetic heterogeneity of cell populations. However, the chromatin accessibility landscape is not well understood and lacks a proper way to interpret it. This work proposes Gene Regulation Accessibility Integrating GeneHancer (GRAIGH), a novel approach to the interpretation of genome accessibility through the integration of the GeneHancer database information, which describes genome-wide enhancer-to-gene associations. Firstly, this paper presents the methods for integrating GeneHancer with scATAC-seq data, creating a new matrix where the features are the GeneHancer elements IDs instead of the accessibility peaks. Secondly, it investigates its capability to analyze the data and detect cellular heterogeneity. In particular, this work shows that the GeneHancer elements are selectively accessible for distinct cell types, and more importantly, their connected genes are precisely known marker genes. Moreover, it investigates the specificity of GeneHancer elements accessibility, demonstrating their high selectivity against the gene activity.

## I. Introduction

Unveiling the complexity of cellular biology is one of the most crucial challenges in biology. Despite the continuous discoveries, many aspects still need to be fully understood [1]. This is especially true for the DNA-related and transcription mechanisms, which are the cardinal stone of cell identities and functionalities [2]. Indeed, gene expression is a crucial part of the homeostasis of a cell and even more essential when looking at multicellular organisms where gene expression profiles affect their distinctive functionality [3]. For this reason, looking at the transcriptomic information of a cell is one of the most effective ways to study the cell type heterogeneity of an organism, to better understand those differences that could be relevant for many different pathologies [4]. The incredible progress in experimental techniques supports this investigation. The Next Generation Sequencing (NGS) technologies permit to perform experiments with a single-cell resolution [5] and a tremendous experimental throughput, allowing simultaneous and fast sequencing of tens to hundreds of cells. Precisely, the well-known and documented single-cell RNA sequencing (scRNA-seq) technology can measure the gene expression profiles of thousands of cells [6] [7] [8]. Despite the incredible value of the data produced with this technology, these data are far from providing a complete view of the biological complexity of cell behavior. Indeed, the transcription of a gene is a convoluted and tightly-regulated process involving many regulatory elements (like promoters, enhancers, transcription factors, etc.) that go beyond mere mRNA synthesis [9] [10]. The interplay of these elements can not be appreciated directly by scRNA-seq data. In order to have a better view of the underlying mechanisms, it is necessary to analyze the genome and, more interestingly, the epigenomic level [11]. Fortunately, new sequencing technologies, such as single-cell assays for transposase-accessible chromatin sequencing (scATAC-seq), allow probing the whole genome chromatin accessibility of single cells. By providing insights into the epigenetic heterogeneity within cell populations, scATAC-seq has opened up new avenues for understanding gene regulation and cellular dynamics [12].

However, the scATAC-seq technology has some limitations regarding the produced data. Differently from scRNA-seq datasets where the features are well-defined set of genes, for scATAC-seq, features are experiment-dependent genomic coordinates [13]. This creates two problems: first, the direct comparison of different datasets is difficult since they may contain different sets of features; second, there is no clear interpretation of what the accessible genomic regions represent. This work proposes Gene Regulation Accessibility Integrating GeneHancer (GRAIGH), a novel computational approach to interpret scATAC-seq features and understand the information they provide. GRAIGH aims to integrate scATAC-seq datasets with the GenHancer database, which describes genome-wide enhancer-to-gene and promoter-to-gene associations. These associations have unique identifiers which have the potential to overcome one of the limitations of the scATAC-seq data, thus enabling interoperability of datasets obtained from different experiments. Moreover, this study shows the strength of employing GenHancer associations to investigate cellular heterogeneity with a higher specificity than other known methods like Gene Activity.

## II. Background

As previously mentioned, scATAC-seq is a powerful technique that allows us to investigate the chromatin accessibility landscape at a single-cell resolution [14]. A dataset obtained from a scATAC-seq experiment is a matrix where each column corresponds to a single cell, and each row is a feature that corresponds to a specific genomic locus. The matrix elements usually provide a binary representation of the chromatin accessibility status for each genomic locus across the individual cells. The features, called peaks, are often associated with regulatory elements, and their presence or absence in a given cell reflects their specific regulatory landscape. However, the definition of the peaks is the outcome of computational algorithms, which take the aligned sequenced fragments (the actual experimental output) and define the peaks as the enriched genomic regions [15]. This process is affected by the algorithm parameters; therefore, the peaks are highly experiment dependent and are not unique and welldefined like genes [16] [17]. This fact limits the power of the scATAC-seq technology since extracting information and creating models from data obtained from different datasets is challenging. Moreover, correlating the trancriptomic and epigenomic levels by linking genes (and their transcription) to specific peaks is not trivial. One of the most known methods to achieve this goal is to employ the concept of Gene Activity Matrix (GAM) [18] [19] [20].

A GAM encodes the genomic accessibility information into a format where genes, rather than genomic regions, serve as features. This representation provides the accessibility level of each gene and its potential for transcription. Many computational methods (e.g., Signac [21] and GeneScoring [22]) define the activity of a gene from the epigenetic signal (i.e., the peaks) overlapping its genomic region and a predefined upstream region of the Transcripton Starting Site (TSS), representing its promoter. These approaches employ the well-documented knowledge about gene body regions inside the genome to interpret and link the peaks to the genes in a trivial way.

However, the peaks overlapping gene bodies and promoters are less than 20% of the total [23], meaning that only looking at them is limiting. With this approach, all remaining epigenetic information is lost. Most of the lost information traces back to imputed enhancer regions [23], but since they are not well-defined, they can not be linked to their affecting genes.

The GenHancer database [24], which provides a database of enhancers and promoter elements linked to their gene of interest, can help address this problem. Therefore, in the following sections, this paper presents a new method, whose workflow is presented in Fig. 1, for integrating GeneHancer with scATAC-seq data. The paper also provides experimental results to show the power of this integration when interpreting and analyzing scATAC-seq data.

**Fig. 1:**
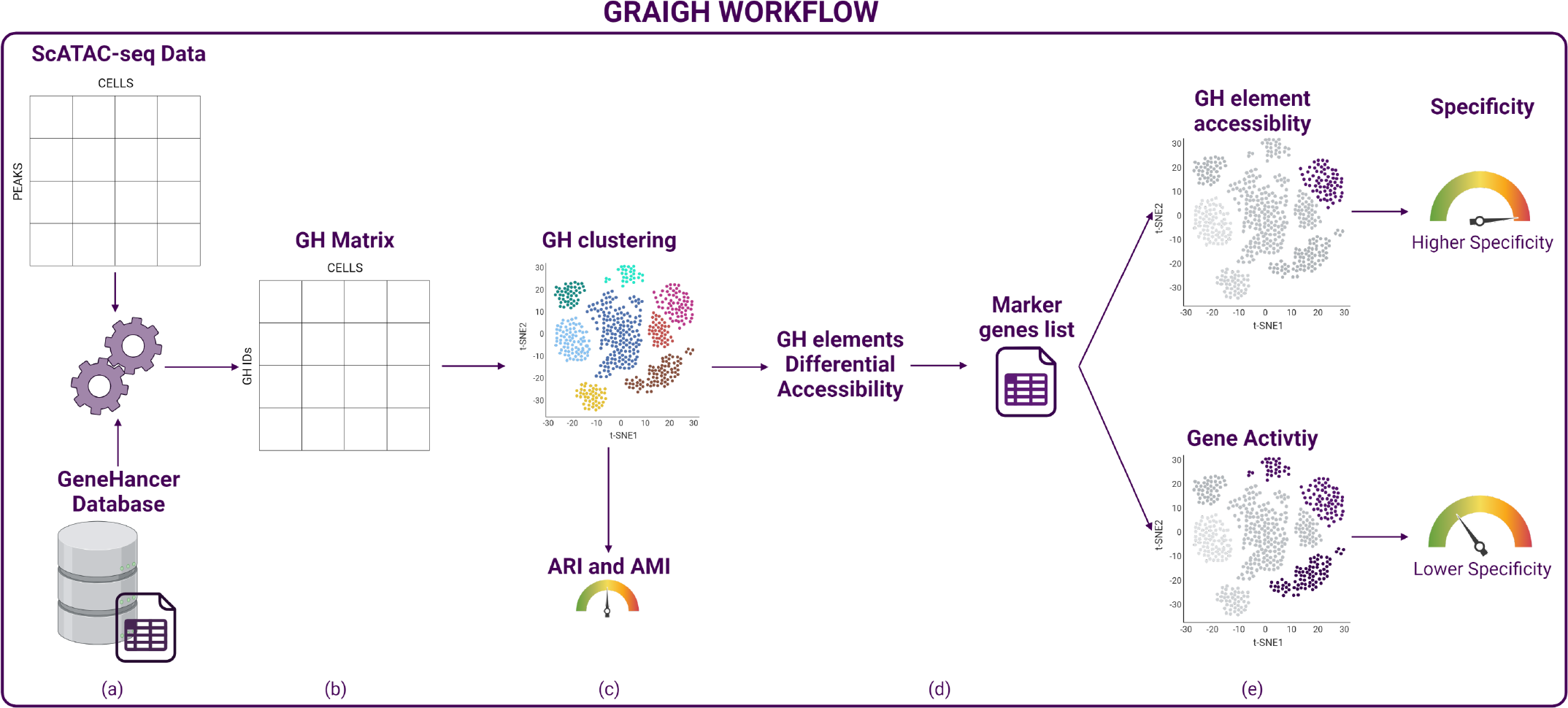
Gene Regulation Accessibility Integrating GeneHancer workflow. (a) The scATAC-seq matrix with peaks as features is integrated with the GeneHancer database. (b) It outputs the GH Matrix with the *GH*_*el*_ IDs as features. (c) The GH Matrix is processed, obtaining the unsupervised clustering. The clustering is compared with the clustering obtained from the processing of the original scATAC-seq data to demonstrate that the GH matrix does not introduce critical biases. (d) Differential Accessibility Analysis of the *GH*_*el*_ returns Enhancers elements related to Known marker genes. (e) Comparison of gene activity and *GH*_*el*_ accessibility of specific marker genes. The results show that the *GH*_*el*_ have higher specificity.

## III. Materials and Methods

### A. Materials

This work employs the latest version of GeneHancer, demonstrating the proposed method on a freely accessible 10xGenomic scATAC-seq dataset, consisting of 10,246 human peripheral blood mononuclear cells (PBMC), with 165,434 called peaks [25]. The code employed in this study is available on GitHub at https://github.com/smilies-polito/GRAIGH. The code is developed using the R language, employing, for the most part, the Seurat suite [26] for the analyses.

### B. GeneHancer

GeneHancer [24] is a part of the well-known GeneCard database [27]. It provides incredible and comprehensive insights into all annotated and predicted human genes. Gene-Hancer is a genome-wide enhancer-to-gene and promoter-to-gene associations database covering up to 18% of the genome. The regulatory elements derive from a cross-source investigation considering up to 9 data sources. It ensures non-redundant, reliable, and comprehensive information on regulatory elements of the human genome with their functional annotation and inferred target genes. Each GeneHancer element (*GH*_*el*_) in the database univocally corresponds to a regulatory element, identified by its genomic coordinates, the genes to which it connects, and a confidence score defining the reliability of the connection. Among the latter, *elite* connections between *GH*_*el*_ and genes are defined as the ones verified by multiple sources. One limitation of the database is that its data are only built for the hg38 genome version. Therefore, it can only support analyses of scATAC-seq data built on the same genome build. The current version of the database comprises 393,464 *GH*_*el*_ and a total of 2,408,198 GH-gene connections, covering 18% of the genome.

### C. scATAC-seq data processing

The workflow to process a scATAC-seq dataset provided in the form of a matrix **A**_|*P* |*×*|*C*|_ (with *P* being the set of peaks and *C* the set of cells) is well-established [28]. This work employs the widely spread R package Seurat to perform data processing.

The processing starts performing the peaks calling step required to identify the available peaks (not required for the employed dataset since peaks are already available). It continues with data normalization and scaling followed by dimensionality reduction using the Latent semantic indexing (LSI) technique. LSI is better suited for this type of data rather than PCA [19]. Then a Uniform Manifold Approximation and Projection (UMAP) [29] (or a Stochastic Neighbor Embedding (tSNE)) dimensionality reduction allows for a two-dimensional dataset representation. From there, unsupervised clustering allows for inspecting the cellular heterogeneity of the dataset. However, this approach still does not provide information on the actual cell types. The Seurat package provides cross-modality data integration and classification to classify the cells with known cell-type labels, employing an external pre-labeled scRNA-seq dataset of the same biological sample. Specifically, the reference dataset employed in this work is a PBMC scRNA-seq dataset suggested by the Seurat guidelines and available at [30]. The results obtained with this classification represent a ground truth useful for further comparisons.

As explained in Section II, a standard tool to elaborate the epigenetic data is a Gene Activity Matrix (GAM). A GAM is a matrix 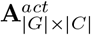 (with *G* being the set of genes and *C* the set of cells) defining the activity as the overall accessibility of a gene. This work employs Seurat’s companion package Signac [21], which provides the procedure for the GAM computation. It is crucial to notice that Signac, as other GAM methods, defines the gene activity from the scATAC-seq signal only from the gene body and its imputed promoter (defined as an upstream region of TSS), so it does not take information from enhancer regions [18] [20].

### D. GH matrix creation and processing

The idea proposed in this paper to integrate GeneHancer with a scATAC-seq dataset is the creation of a new matrix where features (rows) represent *GH*_*el*_ and columns represent cells. The process to create this matrix is formalized in Algorithm 1. The algorithm receives the list of *GH*_*el*_ (**GH**_**el**_ = *{GH*_*el*1_, *GH*_*el*2_, …, *GH*_*elN*_ *}*) and the set of peaks (*P* = *{p*_1_, *p*_2_, …, *p*_*N*_ *}*) genomic coordinates and creates an association matrix 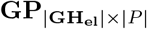 such that each element is one if a peak overlaps a *GH*_*el*_. In the algorithm, the operator *a* ⊆ *b* is used to denote that the two regions *a* and *b* belong to the same chromosome with *a* overlapping *b* (i.e., *a*_*start*_ ≥ *b*_*start*_ ∧ *a*_*end*_ ≤ *b*_*end*_). This process translates the epigenetic features (i.e., peaks) into something univocally defined and comparable with other experiments (i.e., *GH*_*el*_). Multiplying the 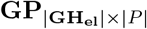 matrix by the **A**_|*P* |*×*|*C*|_ matrix produces the 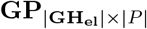 matrix, which provides a new representation of the original scATAC-seq data and can be processed with the same workflow exposed in Section III-C. This procedure leads to a new 2D visualization and clustering of the cells. Comparing the clustering obtained with the two matrices is relevant to prove that the analysis of scATAC-seq datasets can be reliably performed on the GH data without losing information, but with the significant benefit of employing uniquely identified and interoperable features (the *GH*_*el*_ elements), which carry meaningful biological insight.

#### Algorithm 1 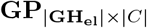 matrix construction

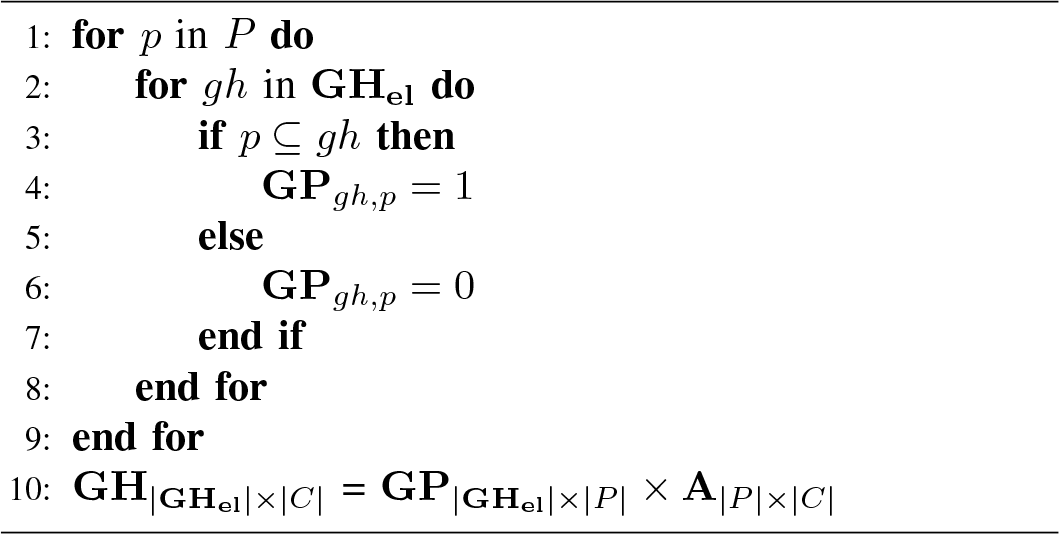

### E. Differentially accessible features and their specificity

Apart from proposing a new scATAC-seq data representation, this paper also aims to investigate if the GeneHancer integration can better support the cellular heterogeneity investigation. Indeed, since the *GH*_*el*_ have defined connections to genes, they could be used to identify cell types in the dataset, similar to the use of gene markers employed when analyzing scRNA-seq data.

The keystone of this approach is the use of differential analysis. Differential analysis is a robust and well-known approach used in single-cell analysis pipelines to study the features that better characterize the different groups of cells, clusters, or cell-type labels. In the case of scRNA-seq data, the Differential Expression (DE) analysis focuses on genes, while for scATAC-seq data, the Differential Accessibility (DA) on the peaks.

Employing the Seurat suite, performing a DA analysis of the *GH*_*el*_ between the cell types is possible. The result is the list of the top *GH*_*el*_ with the highest average log-fold change for each cell type. For each element, the procedure retrieves all the genes with a regulatory link to them if the connection is labeled as elite. Therefore, this work checks the presence of known marker genes from the retrieved list of genes. This indicates how DA *GH*_*el*_ are coherent with specific cell types and can bring up relevant information about the cell heterogeneity of the dataset.

Furthermore, to provide a more comprehensive characterization of both the *GH*_*el*_ and the cell types present, this study introduces an additional quantitative analysis to explore their interrelation.

When dealing with single-cell RNA sequencing (scRNA-seq) data, a common method for deciphering the diversity of cell populations within a dataset involves examining the expression patterns of well-established marker genes. These markers serve to distinguish distinct clusters as specific cell types. Given this, carrying out a similar investigation using the GH matrix becomes intriguing.

Using an approach akin to the one described in the preceding section, this research compiles a list of established marker genes and identifies the associated *GH*_*el*_ with strong connections. These *GH*_*el*_ entities are expected to be uniquely accessible to the same cell type, thereby holding potential as epigenetic markers across various experimental setups.

However, it could be argued that the Gene Activity Matrix (GAM) accomplishes a comparable inquiry with fewer steps by directly assessing the activity of marker genes. Consequently, this study demonstrates that employing the *GH*_*el*_-based approach yields superior outcomes in recognizing distinct cell types than relying solely on gene activity evaluations.

Indeed, one relevant characteristic of a cell type marker for heterogeneity investigation is its specificity [31] defined as:

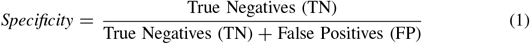

The concept of “high specificity” refers to a characteristic where a feature utilized to distinguish a particular cell type is exclusively present in cells belonging to that specific type. This study assesses the level of specificity in terms of both the accessibility of *GH*_*el*_ and the activity of literature marker genes for various cell types derived from the Seurat integration, subsequently drawing comparisons between them.

When dealing with a marker gene, the initial step involves selecting the *GH*_*el*_ associated with that particular gene. Since a single gene can be linked to multiple *GH*_*el*_, this research focuses on the elite *GH*_*el*_ generated through DE analysis, a collection referred to as 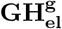. Subsequently, following the outlined Algorithm 2, for a given marker gene *g* corresponding to a specific cell type *t*, the activity vector of *g* denoted as **A**_*g×*|*C*|_, the vector of cell-type labels *CT*, and the accessibility vector of each associated *GH*_*el*_, denoted as **GH**_*gh×*|*C*|_, are binarized (in lines 2 to 7, 10 to 15, and 18 to 23 respectively). With the cell-type labels vector *CT* serving as the reference truth and the activity vector as predictive data, the study calculates the specificity pertaining to the activity (line 16). Subsequently, for each element *gh* within the 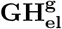 list, the algorithm computes its specificity (line 25), thereby establishing the accessibility specificity as the average value across all elements (line 27).

In conclusion, the algorithm computes the disparity between the specificity concerning *GH*_*el*_ accessibility and activity (line 28).

## IV. Results

### A. GH matrix is equivalent to original data

This section presents results obtained from the application of the proposed methods to the previously mentioned PBMC dataset consisting of 165,376 peaks per 10,246 cells. The aim of these results is to show that a GH matrix is equivalent to the original data and can be used for cell heterogeneity studies with the advantage of easy comparison among multiple datasets.

Fig. 3a shows the cell clustering performed on the original data. Cells are grouped into two significant populations and some smaller groups, which is in line with what is expected for this type of sample [19], divided into 20 clusters by

#### Algorithm 2 Specificity Analysis

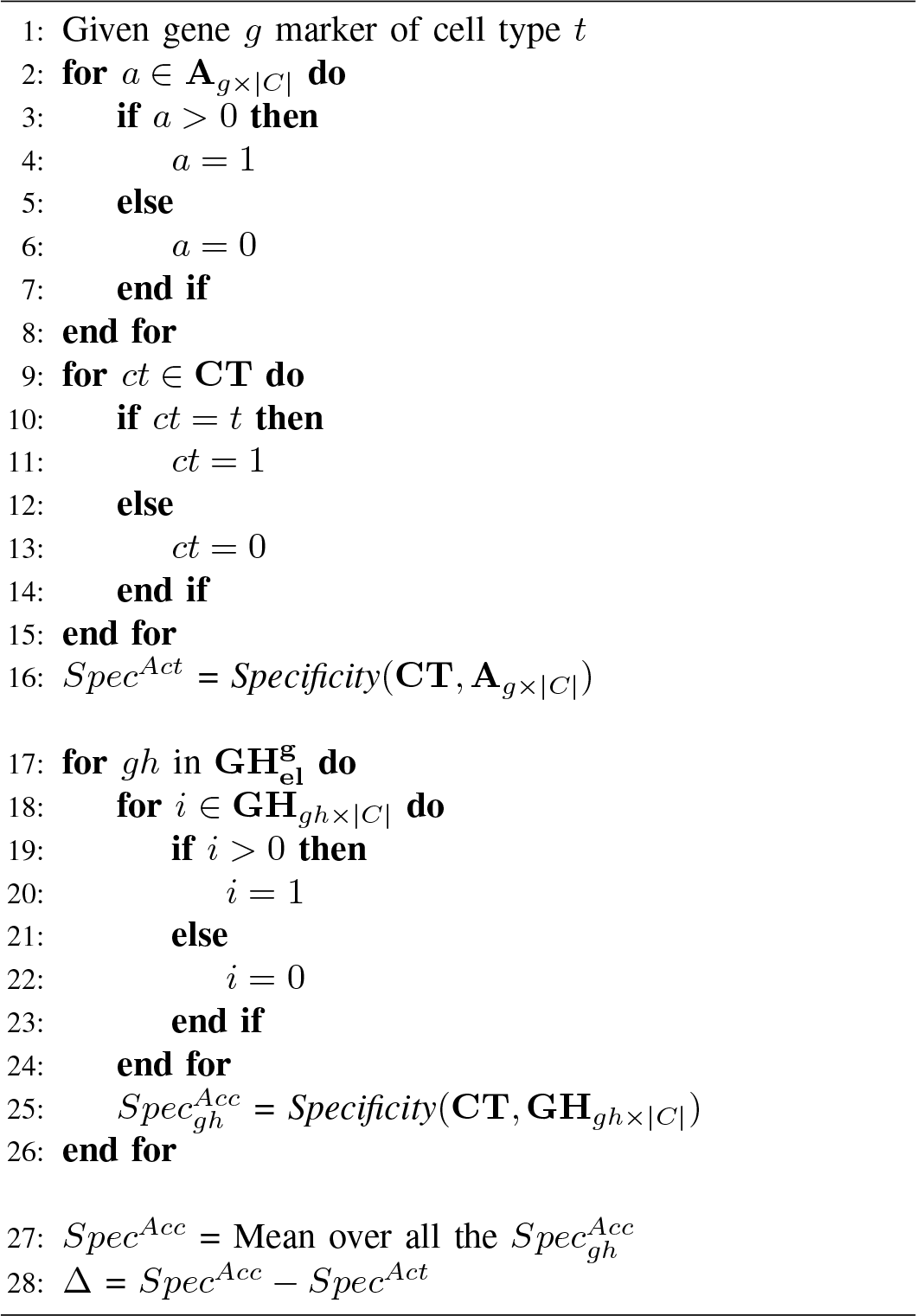

the unsupervised clusterization algorithm. This result is also corroborated by the cell-type classification obtained with the Seurat integration shown in Fig. 3b. The algorithm identifies the T cells (subdivided into subtypes) as the major population, followed by the Monocytes. The two smaller groups represent the Natural Killer (NK) cells and the B cells, ending with a few cells labeled Dendritic cells.

The proposed approach takes the coordinates of the 393,464 *GH*_*el*_ and overlaps them with the coordinates of the peaks, creating the connection matrix. Fig. 2 shows the distribution of how many peaks are connected to the *GH*_*el*_. Most *GH*_*el*_ have one or few peaks overlapping them, with a reduced number overlapping many peaks (up to 27). The reason for so many overlaps stems from the length of the peaks being much smaller than the *GH*_*el*_. After filtering out the *GH*_*el*_ with no overlaps we obtained a 109,620x165,376 connection matrix that, multiplied by the original data, generated a 109,620x10,246 GH matrix.

**Fig. 2:**
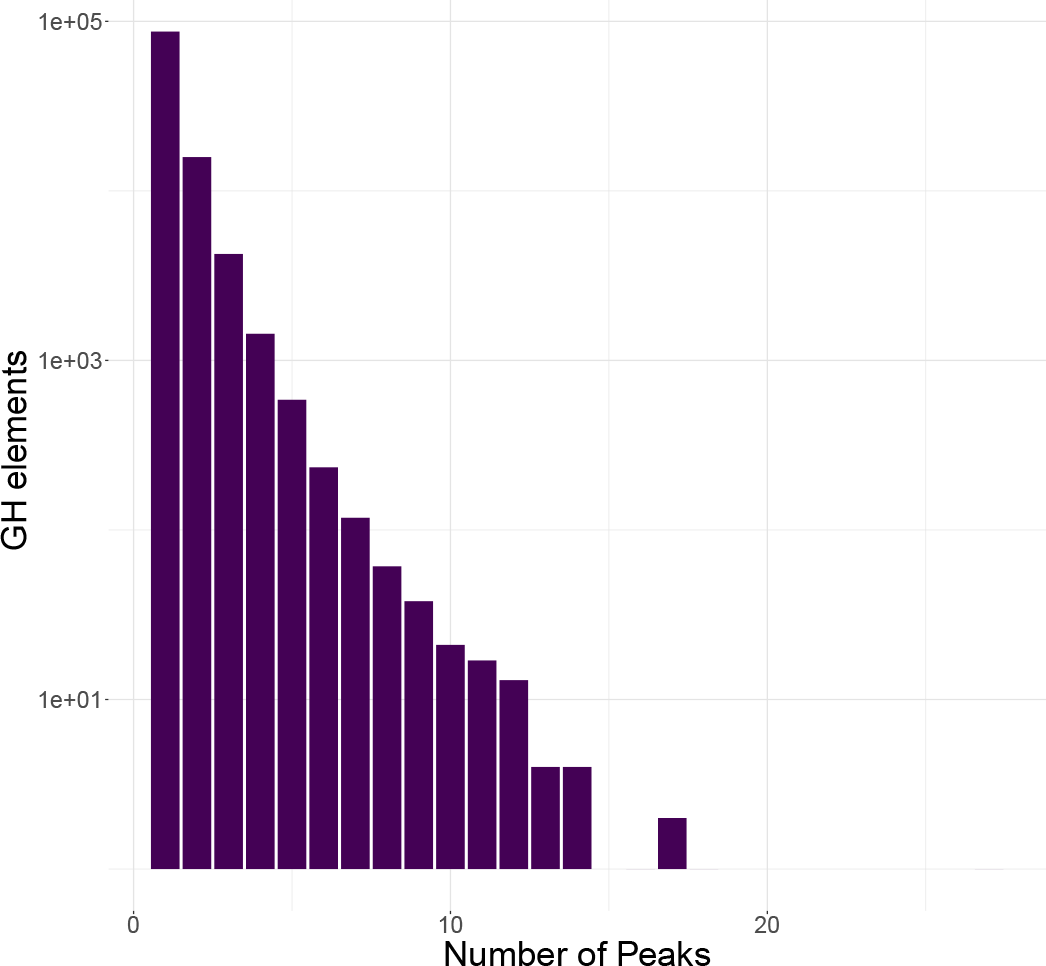
Barplot of Peaks per *GH*_*el*_. The y axis is in logarithmic scale. As the plot shows, the great majority of *GH*_*el*_ have one or few peaks overlapping them. However, in many cases, the *GH*_*el*_ have many peaks overlapping them since they are longer than the peaks.

**Fig. 3:**
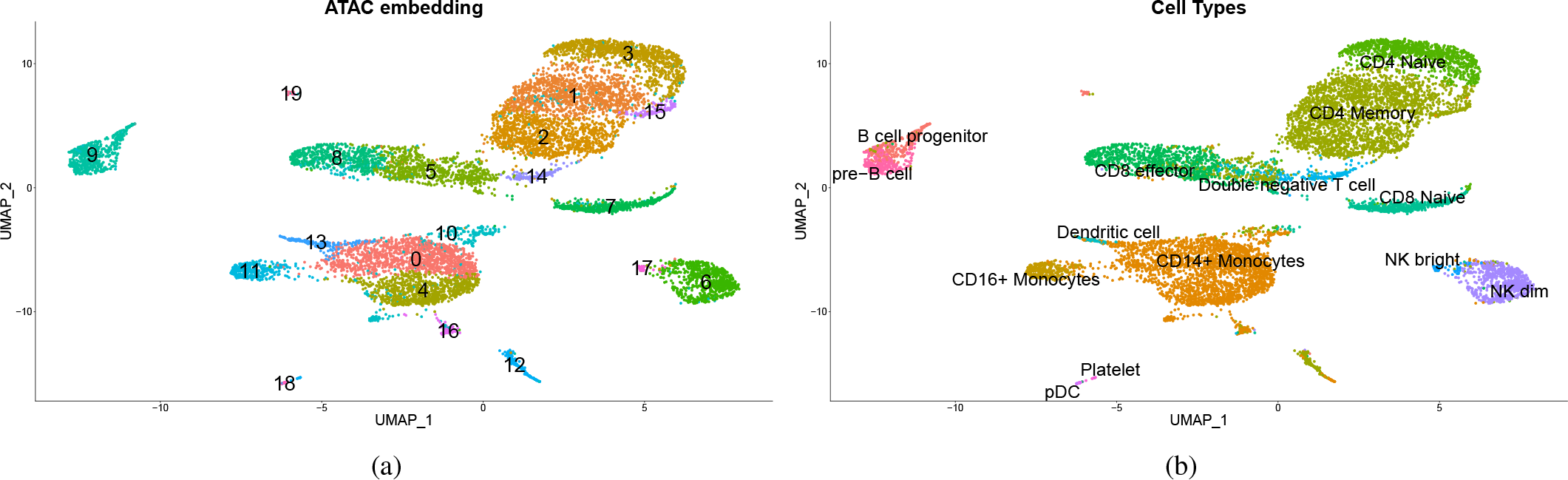
(a) UMAP visualization obtained from processing the original scATAC-seq data, with its unsupervised clusterization. (b) Same UMAP embedding but colored with the cell-type labels obtained from the Seurat label transfer integration.

Fig. 4 displays a 2D representation of the data obtained from processing the GH matrix, showcasing cluster patterns similar to those illustrated in Fig. 3a. In this instance, the unsupervised clustering algorithm has identified 19 clusters, and the evident resemblance among the identified clusters can be easily discerned. Additionally, it is feasible to assess their similarity using metrics such as Adjust Rand Index (ARI) and Adjust Mutual Information (AMI), commonly employed to gauge classification similarities [19] [31].

**Fig. 4:**
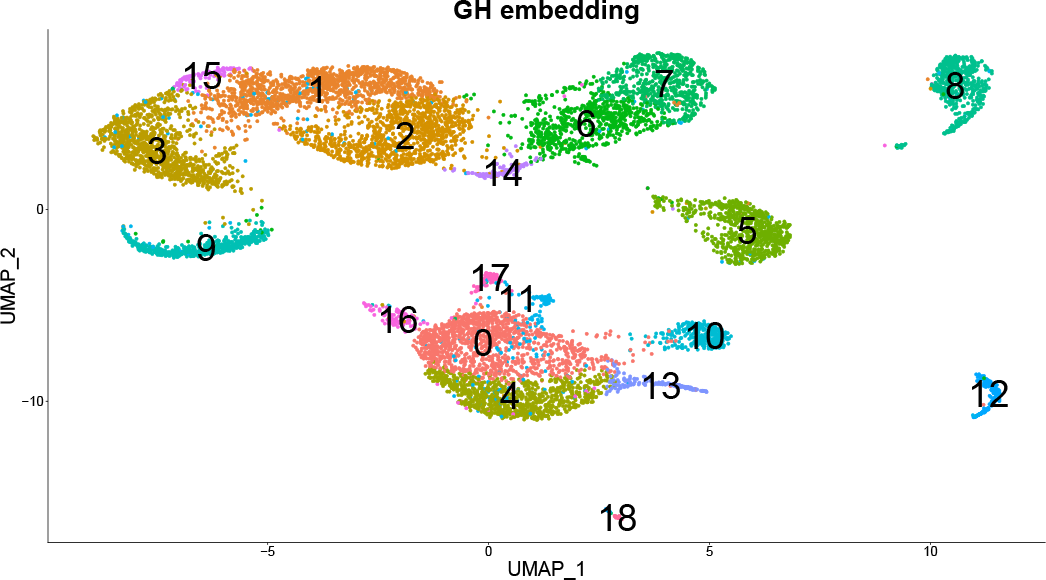
UMAP embedding and clustering result, obtained from the processing of the GH matrix. Comparing with Fig. 3.a, it is evident that the clustering subdivision of the cells is mostly coherent, demonstrating GH matrix does not introduce relevant biases and can be used to process scATAC-seq data.

When these metrics are computed for the two clustering results, they yield an AMI of 0.867 and an ARI of 0.804, underscoring a significant likeness between the two clusterings. This outcome underscores the credibility of the GH matrix approach in producing comparable outcomes to the original data while avoiding noticeable biases. Consequently, it confirms the effectiveness of this approach, which leverages biologically meaningful features to not only appropriately analyze scATAC-seq data but also to directly compare results across different experiments.

### B. GH_el_ accessibility are cell-type specific

The outcomes presented in this section are geared towards exploring the effectiveness of the *GH*_*el*_, and in a broader sense, the utility of this technique in conducting an analysis of cell heterogeneity. Additionally, these results are compared to those of the Gene Activity Matrix. In the course of processing the initial dataset, a GAM comprising 19,607 genes is produced. This particular GAM serves as the benchmark against which the final analysis is evaluated.

The DA analysis on the GH matrix identifies 76,081 imputed DA *GH*_*el*_, of which 73,235 are positive markers, meaning they positively characterize the corresponding cell type.

Investigating some of them brings out some interesting *GH*_*el*_. For example, the elements GH02J086783 and GH02J086805, specific for the CD8 subtypes cells, are elite enhancers for CD8A and CD8B genes, known markers of the homonymous cells. The same reasoning can be done for GH05J140611 and GH05J140596, differentially accessible for the Monocytes and elite enhancers of the CD14 gene. These results show that the DA *GH*_*el*_ are selectively accessible for distinct cell types. Furthermore, genes connected to them are markers of the distinct cell types. This highlights the coherence of accessibility of regulatory elements and known cell-types and can bring crucial information for cellular heterogeneity investigation.

Finally, this study investigates whether *GH*_*el*_ connected to cell-type marker genes have a valuable discriminating power in analyzing cell heterogeneity. In particular, it compares the specificity of the accessibility of those *GH*_*el*_ versus the specificity of the activity of their target marker genes. Table I reports such specificity values for the list of considered marker genes and their difference.

**TABLE I:**
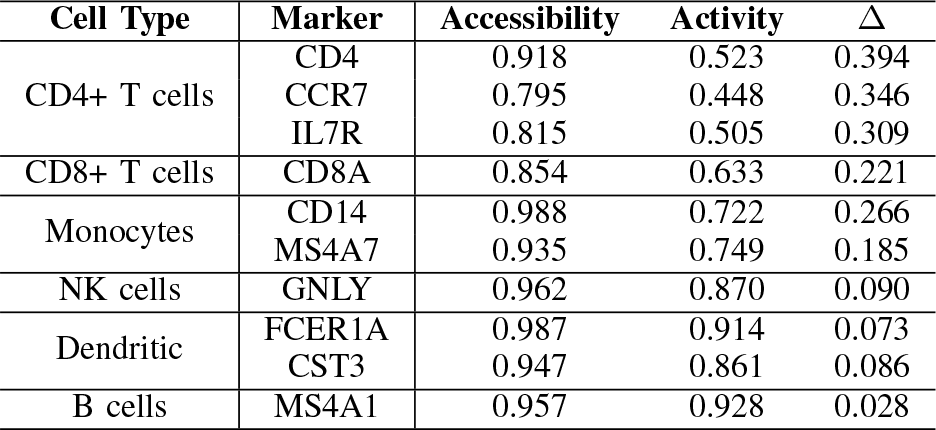
The table reports for each main cell type and marker gene the specificity of the gene activity and the mean of the *GH*_*el*_ accessibility. The last row is the difference which highlights how the *GH*_*el*_ accessibility has always higher specificity.

The reported results are intriguing. First, all the differences are positive, meaning the *GH*_*el*_ specificity is always higher than the activity specificity. Moreover, the differences are particularly significant, especially for more populated cell types like T cell subtypes and Monocytes, which go up to 0.266 and 0.394, respectively. The difference is lower for smaller populations, like Dendritic and B cells, since well-separated subtypes tend to be also well-defined at the activity level. However, their *GH*_*el*_ specificity remains consistently higher than 0.9 showing the method reliability. It becomes even more evident when plotting the features on the dataset. Fig. 5 shows both the accessibility of the GH05J140611 (a), an enhancer element of the gene CD14 marker of Monocytes, and the same gene’s activity (b). One immediately can notice how the gene activity spreads all over the dataset, while the accessibility of its enhancer element is specific for the Monocytes population. Similarly, Fig. 6 shows the same representation for the CD4 gene and the accessibility of the connected element GH12J006784. Even in this case, it is clear how the accessibility of the *GH*_*el*_ is more specific than the activity.

**Fig. 5:**
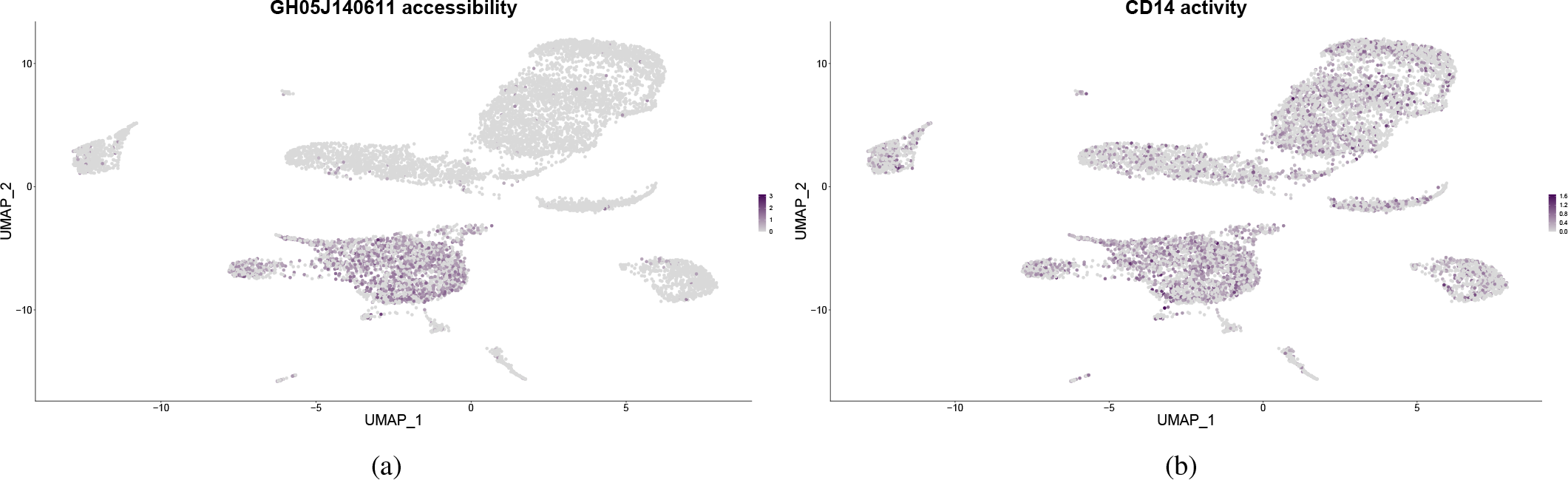
The CD14 gene is a marker for the Monocytes. The accessibility of its enhancer GH05J140611 is specifically accessibile in the Monocytes population (a), while its gene activity (b) is more spread out in many other cells.

**Fig. 6:**
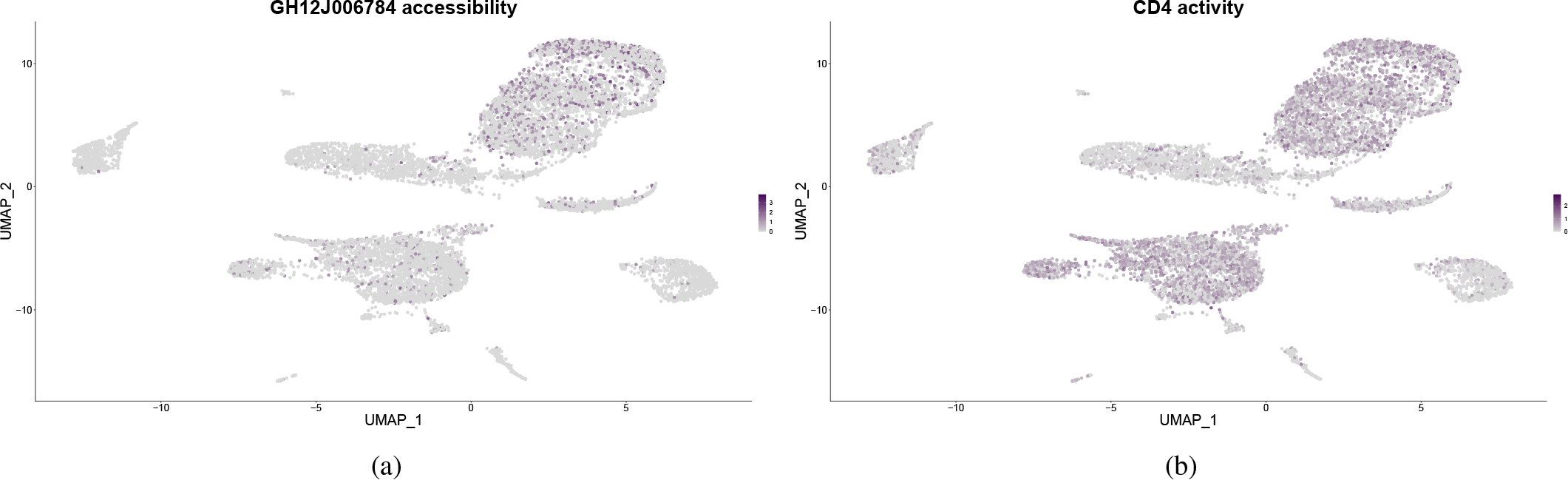
The CD4 gene is a marker for the homonymous T cells. Its enhancer GH12J006784 is specifically accessibile in the CD4+ T cells population (a), while its gene activity (b) is more spread out in many other cells.

These results show how the use of *GH*_*el*_ as features is a reliable way to interpret the scATAC-seq data, and, more importantly, they have greater specificity in detecting the heterogeneity of the dataset than the gene activity.

## V. Conclusions

In conclusion, this work proposes a novel approach, Gene Regulation Accessibility Integrating GeneHancer (GRAIGH), to interpret and understand scATAC-seq data. The scATAC-seq data, which provides information about the chromatin accessibility landscape at single-cell resolution, has been challenging to interpret due to the lack of well-defined features like genes in scRNA-seq data. The paper introduces the integration of the scATAC-seq data with the GeneHancer database, which describes genome-wide enhancer-to-gene and promoter-to-gene associations. The *GH*_*el*_ have the robust quality of having unique identifiers, allowing for the interoperability of results between different datasets.

This study demonstrates that integrating GeneHancer data with scATAC-seq and employing the *GH*_*el*_ as features instead of the peaks is a valid and powerful way to investigate single-cell epigennomic data. The approach is validated by comparing the results obtained from the GH matrix data with the original scATAC-seq data, showing the integration does not introduce any significant biases.

Moreover, the paper shows that the accessibility of *GH*_*el*_ elements pertains to particular cell types. By analyzing the *GH*_*el*_ accessibility of known marker genes, the study demonstrates that *GH*_*el*_-based analysis provides higher specificity than traditional gene activity analysis in identifying cell types.

Certainly, this approach comes with certain limitations. First and foremost, the availability of GeneHancer information is restricted solely to the human genome. Moreover, understanding transcriptomic regulation involves navigating a complex and constantly changing process, necessitating comprehensive information from various sources to achieve a thorough comprehension. In light of this, forthcoming advancements will strive to incorporate motif information from the peaks, a critical component for discerning the Transcription Factors contributing to regulation. Furthermore, an enticing prospect lies in employing multi-omic datasets to explore the correlation between *GH*_*el*_ accessibility and actual gene expression.

Overall, the GRAIGH approach presents a valuable means of interpreting scATAC-seq data, offering insights into the intricate regulatory mechanisms that underscore cellular heterogeneity. This method opens up novel avenues for delving into gene regulation and cellular dynamics, carrying significant implications for comprehending cell characteristics, functions, and their relevance to various medical conditions. In conclusion, integrating scATAC-seq data with the GeneHancer database represents a promising stride towards unraveling the complexities of cellular biology at the epigenomic level.

## VI. Data and code availability

GeneCard allows direct download of the older database 2017 version https://www.genecards.org/ GeneHancer Version 4-4, but it is possible to request the authors’ access to the latest versions from their online platform https://www.genecards.org/Guide/DatasetRequest. The 10X genomics dataset is freely available at https://www.10xgenomics.com/resources/datasets/ 10k-human-pbmcs-atac-v2-chromium-controller-2-standard. All the code employed in this study is publicly available on the GitHub repository at https://github.com/smilies-polito/GRAIGH.

